# SPCoral: diagonal integration of spatial multi-omics across diverse modalities and technologies

**DOI:** 10.64898/2026.02.02.703207

**Authors:** Heqi Wang, Jiao Yuan, Kunshi Li, Xufeng Chen, Xi Yan, Ping Lin, Zhixuan Tang, Baihua Wu, Haitao Nan, Yiwei Lai, Yuan Lv, Miguel A. Esteban, Lu Xie, Gangqi Wang, Lijian Hui, Hong Li

## Abstract

Spatial multi-omics is indispensable for decoding the comprehensive molecular landscape of biological systems. However, the integration of multi-omics remains largely unresolved due to inherent disparities in molecular features, spatial morphology, and resolution. Here we developed SPCoral for diagonal integration of spatial multiomics across adjacent slices. SPCoral extracts spatial covariation patterns via graph attention networks, followed by the use of optimal transport to identify high-confidence anchors in an unsupervised, feature-independent manner. SPCoral utilizes a cross-modality attention network to enable seamless cross-resolution feature integration alongside robust cross-omics prediction. Comprehensive benchmarking demonstrates SPCoral’s superior performance across different technologies, modalities and varied resolutions. The integrated multi-omics representation further improves spatial domain identification, effectively augments experimental data, enables cross-modal association analysis, and facilitates cell-cell communication. SPCoral exhibits good scalability with data size, reveals biological insights that are not attainable using a single modality. In summary, SPCoral offers a powerful framework for spatial multi-omics integration across various technologies and biological scenarios.

## Introduction

Over the past decades, spatial omics technologies have achieved tremendous advances in both sequencing throughput and spatial resolution^1,2^, and are being applied to various biology researches^3,4^. Spatial transcriptomics, such as 10x Xenium^5^, Stereo-seq^6^, and CosMx^7^ has elevated the resolution of captured RNA to single-cell or even sub-cellular levels. Spatial proteomics technologies such as CODEX^8^ and IMC^9^ enable the simultaneous detection of expression signals from dozens of proteins. Spatial metabolomics technologies like MALDI-2^10^ can also achieve untargeted metabolite profiling through spatially resolved mass spectrometry. Given the complementarity of different omics approaches, a growing number of researchers are attempting to measure and integrate different omics data^11^. However, currently, only a few experimental techniques can simultaneously measure two types of omics data from the same tissue slice^12^. In most cases, two types of omics data were measured from adjacent slices by independent technologies. This gave rise to a major challenge: how to perform diagonal integration of distinct spatial omics data measured from two adjacent tissue slices— that is, simultaneously achieving alignment of slices without shared features and integration of information from spatial data at different resolutions^13,14^.

Several published methods, such as scSLAT^15^, SANTO^16^, and CAST^17^, employ graph neural network models for adjacent slice alignment. These approaches enable cross-resolution data registration but reply on shared features. Despite claims of supporting cross-omics alignment, most tools still require pre-established correspondences between multi-omics features. For example, converting proteins or ATAC-seq peaks to genes. This task is infeasible for some omics data without clear interconnections, such as transcriptomes and metabolomes. Another category of methods aims to jointly embed two omics datasets obtained from the same spatial locations into a shared latent space. This includes early work such as WNN^18^, which integrates single-cell multi-omics data, as well as more recent methods like SpatialGlue^19^, MISO^20^, and SMOPCA^21^, which are specially tailored for spatially resolved multi-omics data. A critical limitation of these approaches is that they require the two omics must be of matching resolution, ensuring a one-to-one correspondence between cells or spots. This requirement is often incompatible with spatial omics data generated by different technologies.

Two newly developed method SpatialMETA^22^ and SpatialEx^23^ leverage histological images as shared features to achieve diagonal integration of multi-omics across slices, but both of them need the pre-registration of histological image with the omics. Additionally, SpatialMETA is designed only for spot-resolution spatial transcriptomics and metabolomics, may not be suitable for other omics or high-resolution technologies; and SpatialEx needs precise cell segmentation and assumes consistent spatial resolution across different measurements.

As far as we known, no existing methods can achieve diagonal integration of spatial multi-omics data from adjacent slices without relying on external information or predefined feature correspondences. To bridge this gap, we present SPCoral (**Sp**atial **C**ross multi-**O**mics **R**egistration and **A**na**L**ysis), which realizes the diagonal integration of different spatial omics from adjacent slices through two dedicated modules: the alignment module combines a graph neural network and fused Gromov-Wasserstein optimal transport^24^ (FGW-OT) to automatically identify paired anchors across two spatial omics; the integration module employs cross-graph learning to perform feature fusion on spatially aligned but modality-different omics data, ultimately yielding an embedding of multi-omics representation. We first tested SPCoral on simulated and experimentally acquired data to benchmark its performance with other methods. Then we then applied SPCoral to integrating spatial transcriptome, epigenome, translatome, proteome, or metabolism data acquired from distinct tissue types. These results highlight SPCoral’s superior performance in aligning and integrating spatial multiomics from adjacent slices, identifying fine-grained spatial domains, cross-modal prediction, and revealing associations among different modalities.

## Results

### Overview of SPCoral

SPCoral was designed to address the challenge of diagonal integration in spatial multi-omics data that generated by different technologies from adjacent tissue slices^25^ (**Fig. 1a**). It consists two core modules: alignment and integration (**Fig. 1b**). Unlike existing methods, the alignment module of SPCoral does not need to rely on shared features between two omics. Instead, SPCoral aims to capture similar “spatial variation patterns” across different omics in adjacent slices. Within a tissue, the complex regulatory networks between omics often make it difficult to establish clear linear correspondences between individual features. However, spatially differential expression, driven by biological processes, may produce comparable patterns of variation across modalities, regardless of the specific features measured. For example, at the interface between tumor and normal tissue, multiple omics (transcriptomics, proteomics, metabolomics, etc.) typically exhibit pronounced differences, even if the underlying regulatory relationships remain unclear or nonlinear. These spatially covarying patterns, rather than the original omics features, serve as shared information across different modalities that SPCoral extracts and leverages to achieve alignment across omics.

**Fig 1.**
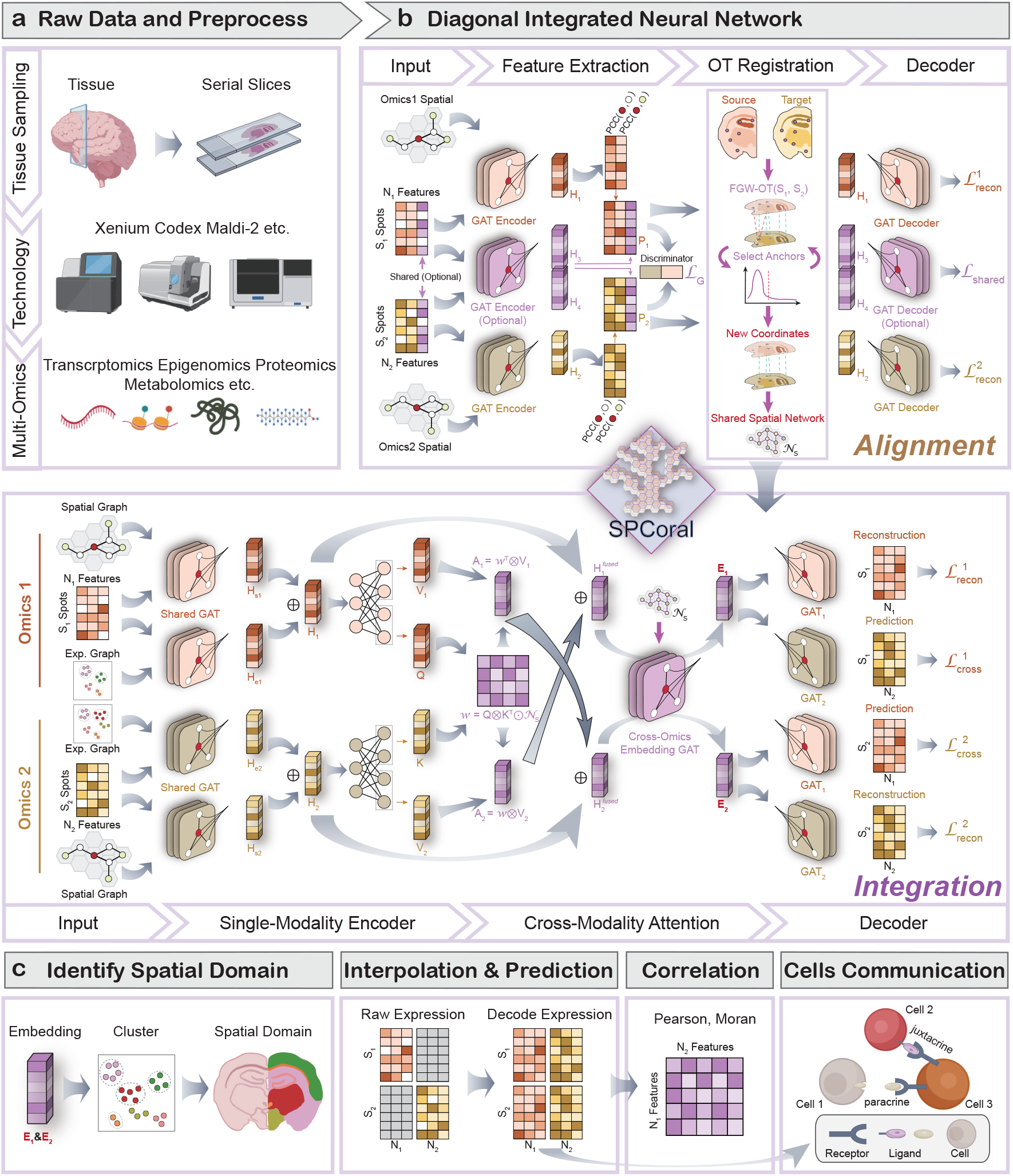
Architecture of the SPCoral framework. **a**, SPCoral is designed to support diagonal integration across various spatial omics modalities from adjacent tissue sections of the same sample. Portions of the schematic diagrams were created with BioGDP.com. **b**, Overview of the SPCoral diagonal integration architecture, which consists of two main components: alignment and integration. In the alignment module, multiple GAT encoders are first employed to perform feature dimensionality reduction on both shared (if available) and modality-specific information for each omics layer. Subsequently, a new shared representation is computed for each spot by calculating PCC with multi-order neighbors. This shared representation is then used in FGW-OT to identify optimal anchor points, achieving spatial alignment between the two slices. In the integration module, each modality is first processed independently using shared-parameter GAT encoders on both spatial and expression-based neighbor graphs for dimensionality reduction. Cross-modality attention learning is then performed based on the reduced features and the cross-omics neighborhood derived from alignment, enabling bidirectional feature fusion to generate joint representations of the two modalities. Finally, GAT-based decoders are applied for feature reconstruction and cross-modality prediction. **c**, Downstream analyses. Embeddings from the Integration module enable spatial domain clustering. Outputs from the decoders facilitate imputation of original expression profiles and cross-modality prediction. The predicted expression profiles support computation of cross-modality correlations and cell–cell communication analysis.

SPCoral first constructs a graph for each omics based on spatial coordinates and learns feature embedding using graph attention networks (GAT). It then computes the Pearson Correlation Coefficient (PCC) between each cell/spot’s embedding and the mean embedding of its multi-order neighbors to capture cross-omics shareable patterns of spatial expression variation. Simultaneously, an optional shared-feature GAT encoder is provided to perform conventional dimensionality reduction and embedding of overlapping features between the two omics. The PCC embeddings are concatenated with the optional shared-feature embeddings to form a joint latent representation for each omics layer. These joint representations are aligned into a common latent space through adversarial training, combined with a decoder-based reconstruction loss. Finally, FGW-OT is applied on the aligned latent space to compute a cross-omics transport plan, from which high-quality anchor pairs are selected to generate a spatial transformation matrix for alignment. This alignment is subsequently leveraged by the integration module for spatial multi-omics feature fusion.

Inspired by cross-graph learning^26^ and multi-modal transformer architectures^27^, the integration module performs cross-modal fusion on spatially aligned multi-omics data to achieve the joint embedding of the two omics. First, within each modality, features are extracted using GAT by jointly considering the spatial proximity and the expression similarity. Subsequently, during the modality fusion stage, unlike conventional integration methods that compute the modality weights only at strictly corresponding positions, SPCoral distributes attentions from each cell/spot to the neighborhood in the adjacent slice. By employing a Query-key-Value mechanism from Transformer^28^, it aggregates complementary information from the other modality. This design enables the modal to produce multi-omics representations even when the two modalities have different resolutions.

Leveraging the resulting multi-modal embedding, users can conduct clustering analysis to define spatial domains that simultaneously reflect information from both omics datasets. Furthermore, through cross-modality decoder architectures, SPCoral can reconstruct imputed expression profiles within each omics dataset and predict the expression of the counterpart omics modality. This not only mitigates the sparsity of spatial data but also improves the downstream analysis, such as spatial correlation of cross-omics features and cell-cell communications. (**Fig. 1c**).

To evaluate the effectiveness of the proposed SPCoral model, we first assessed the alignment and integration modules separately on multiple simulated datasets and benchmarked them against the state-of-the-art methods in their respective domains. Then we applied SPCoral to five experimental datasets, conducting comprehensive method validation and downstream task evaluation to highlight the distinctive advantages of SPCoral.

### Systematic benchmark suggests SPCoral is accurate and robust for multi-omics alignment

First, we utilized scMutiSim^29^ to generate simulated spatial ATAC-seq and RNA-seq datasets comprising four distinct spatial layouts to evaluate SPCoral’s alignment capability across diverse tissue architectures (**Fig. 2a**). We compared SPCoral with existing alignment tools (scSLAT, SANTO, and CAST) using the *Norm Error*, defined as the average distance between the aligned spots and their respective landmarks. Across all four simulated layouts, SPCoral achieved the best overall registration performance (**Fig. 2b**). Furthermore, we conducted additional tests on other randomly generated samples for each of the four layouts (**Extended Data Fig. 1**). The results demonstrate that SPCoral exhibits stable performance when faced with varying spatial structures.

**Fig 2.**
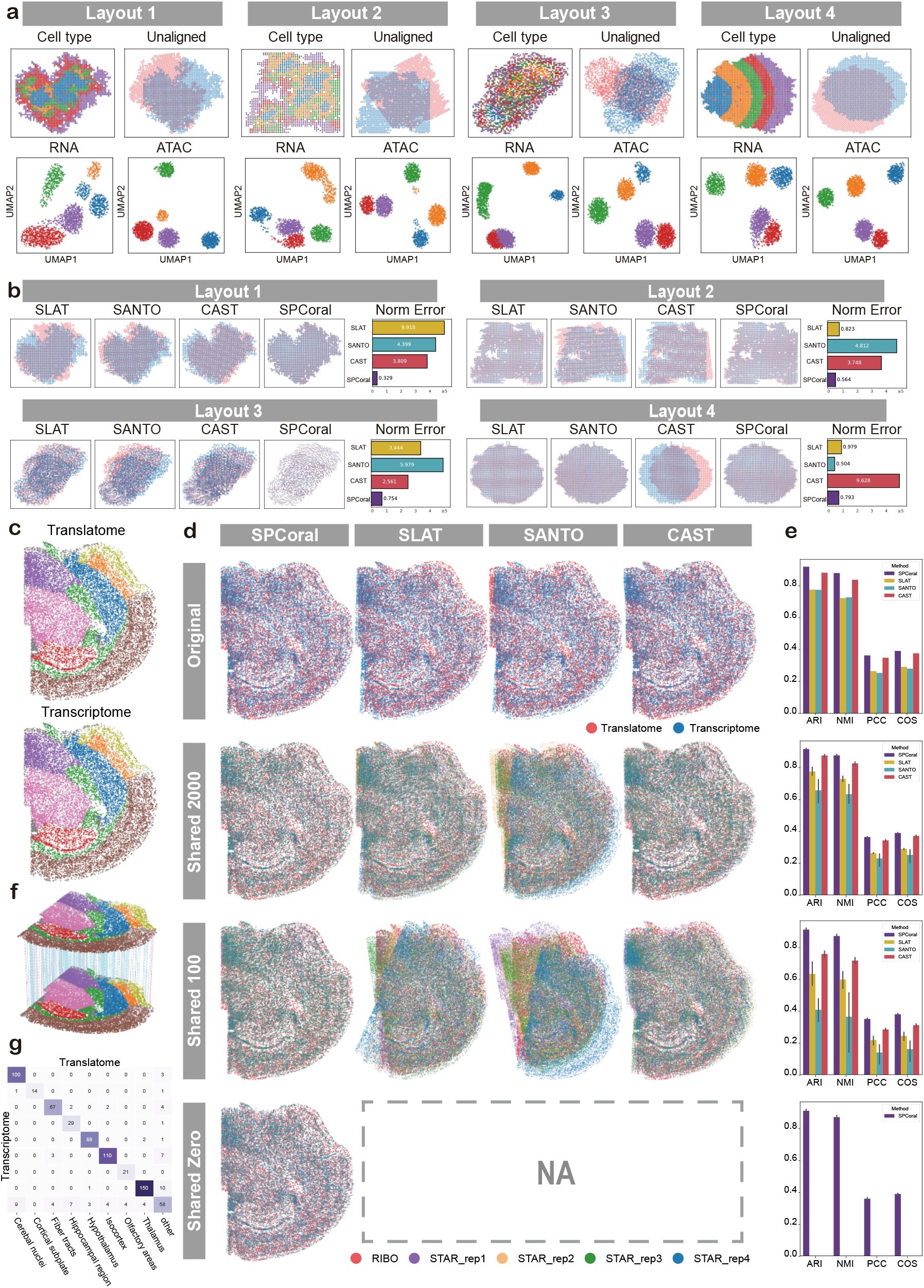
Benchmarking of the SPCoral alignment module on simulated and real data. **a**, Description of simulated data. Four spatial multi-omics datasets with different layouts were generated using scMultiSim. Each dataset contains five virtual cell types along with simulated RNA and ATAC expression profiles. Unaligned data were constructed by randomly shuffling the spatial coordinates of the RNA modality. **b**, Performance of four methods (SLAT, SANTO, CAST, and SPCoral) on the four simulated datasets. Bar plots show the mean error (distance between aligned spot coordinates and ground truth) for each method, with the x-axis capped at 5. **c**, Spatial transcriptomics (STARmap) and translatomics (RIBOmap) data of mouse brain, colored by the annotated tissue regions provided in the original paper. **d**, Alignment results of the four methods on the mouse brain dataset, with translatomics as the target and transcriptomics as the source. From top to bottom: original dataset, subsampled dataset with 2000 shared genes, subsampled dataset with 100 shared genes, and subsampled dataset with no shared genes. Each subsampling process was repeated five times. **e**, Quantitative evaluation of alignment performance using four metrics. Each subplot corresponds to the subplot in **d**, with error bars indicating the range across the five random subsampling replicates. **f**, Visualization of 300 randomly selected anchor points. The upper is the translatomics, and the lower is the transcriptomics. Blue lines indicate that anchors are located in the same tissue regions of the two omics datasets; red lines indicate that anchors are in mismatched regions. **g**, Statistical summary of region correspondence between paired anchors. Row names and column names follow the same order.

Since fully simulated data cannot completely recapitulate the complexity of real tissues, we next evaluated SPCoral on experimental datasets. Here we utilized spatial transcriptomics (STARmap) and translatomics (RIBOmap) data acquired from adjacent tissue slices of mouse brain^30^ (**Fig. 2c**). To explicitly demonstrate SPCoral’s unique advantage in handling spatial multi-omics data in minimal or no shared features, we constructed shared features by randomly subsampling 2000, 100, or 0 genes from the total 5413 genes, while the remaining genes were randomly assigned to only one modality. The subsampling process were repeated five times by different random seeds. SPCoral and the three competing methods (scSLAT^15^, SANTO^16^, and CAST^17^) were evaluated on these datasets using four metrics (**Fig. 2d**, full experimental details are in the **Supplementary Information**). SPCoral achieved the best performance on the original dataset and crucially maintained stability as the number of shared features decreased. In contrast, all competing methods exhibited marked performance degradation with fewer shared features. More importantly, SPCoral is the only method that can align two omics when no features were shared. These results highlight SPCoral’s robustness and its ability to align adjacent slices reliably under realistic conditions where shared features are scarce or absent.

To further illustrate the anchor refinement progress during SPCoral’s alignment, we visualized the extracted anchor pairs and counted the number of anchors that belong to the same tissue regions in the two omics datasets. After five iterative rounds of filtering, incorrectly matched anchors were effectively removed, resulting in a marked improvement in overall alignment performance (**Extended Data Fig. 2a-c**). The final alignment used the anchors obtained through ten iterative rounds (**Fig. 2f, g**). For most regions, around 99% of the anchors were correctly matched; even for the most challenging ‘Fiber tracts’, the correcting matching rate still reached 96%. These results demonstrate the effectiveness of SPCoral’s anchor refinement procedure.

### SPCoral enhances spatial multi-omics integration and spatial architecture delineation

After coordinate alignment, the integration module learns a low-dimensional integrated embedding from two data modality, and then we can use clustering to identify spatial domains. To benchmark SpatialGlue’s integration module, we first compared SPCoral with four competing methods (Seurat_WNN^18^, SpatialGlue^19^, MISO^20^, and SMOPCA^21^) using the simulated spatial transcriptome and proteome data generated by SpatialGlue. Specifically, we constructed three independent simulated datasets with ground-truth spatial domains (factors) for each modality. For each dataset, we generated two versions of modality 2 data: one with resolution identical to that of modality 1 and the other with the mismatched resolution (**Fig. 3a**). Visually, the clustering of embedding from spatial-aware methods substantially outperforms those from the non-spatial Seurat_WNN approach. SPCoral successfully recovered all five spatial factors, and maintained excellent performance even when resolutions differed between two modalities (**Fig. 3b**). In the UMAP visualizations of integrated embeddings, spots from the two mismatched-resolution modalities were seamlessly fused into unified clusters, with all five factors clearly distinguishable (**Fig. 3c**). Quantitatively, with same-resolution data, SPCoral outperformed all competing methods across all six metrics. With mismatched-resolution data, SPCoral exhibited no substantial performance drop (**Fig. 3d**). Across all three independence datasets, SPCoral performed the best with little variance and demonstrated strong robustness (**Fig. 3e, Extended Data Fig. 3**).

**Fig 3.**
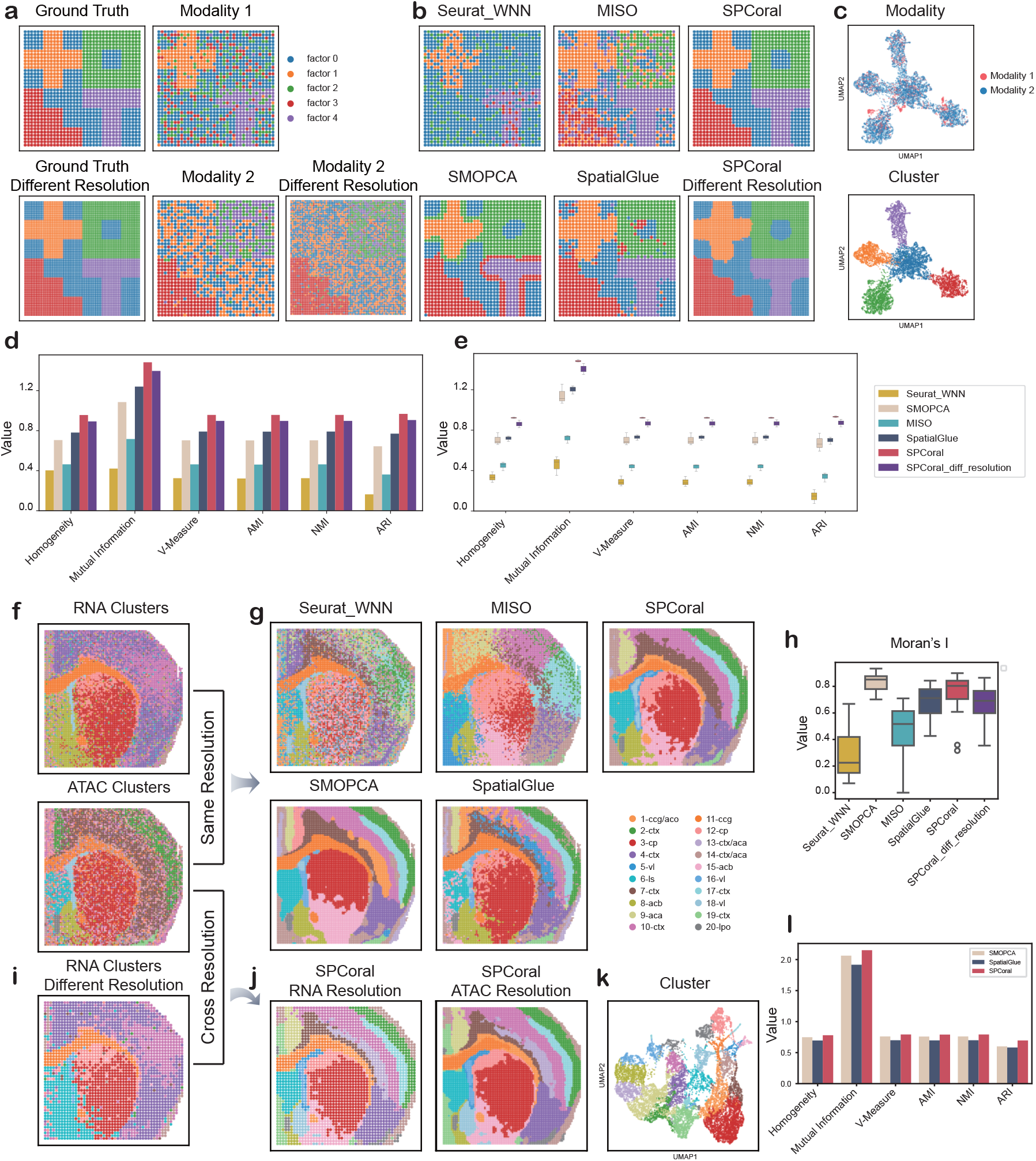
Evaluation of the integration module of SPCoral on simulated and real data. **a**, Simulated data used for evaluating the integration model. From top to bottom and left to right: ground-truth spatial domains at default resolution (1296 spots, 36 × 36 grid), ground-truth spatial domains at higher resolution (5184 spots, 72 × 72 grid), clustering results for modality 1 at default resolution, clustering results for modality 2 at default resolution, and clustering results for modality 2 at higher resolution. Colors represent the spatial factors that generated the simulated data. **b**, Multi-modal integration results from SPCoral and four competing methods (Seurat WNN, SMOPCA, MISO, and SpatialGlue). SPCoral results are shown for both matched-resolution and mismatched-resolution data. **c**, UMAP visualizations of SPCoral embeddings for the cross-resolution data, colored by modality or by spatial domain labels. **d**, Quantitative evaluation of the methods in the simulated dataset (**a)**. **e**, Box plots of the six metrics across the three simulated datasets. The box limits denote the upper and lower quartiles. **f**, The spatial transcriptomics-epigenomics dataset of a mouse brain. The clustering results of each single omics were obtained from the original paper. **g**, In situ spatial domain maps derived from multi-omics embedding clustering. Clustering was performed with reference to the annotations provided in the original SpatialGlue study. The color scheme is defined based on SPCoral results; annotations for other methods are aligned with SPCoral as closely as possible, though they may not always match the same anatomical structures. **h**, Spatial continuity scores for the domains, measured by the Moran’s index. **i**, Clustering results of the downsampled transcriptome data. **j**, Clustering results from the cross-resolution integration of SPCoral. Clusters are displayed on the in-situ images of two slices. **k**, UMAP visualization of the cross-resolution integration results. **l**, Quantitative evaluation of the cross-resolution integration. Spatial domains derived from the original data using three top-performing methods were quantitatively compared with those obtained from cross-resolution data via SPCoral.

Next, we used a real spatial transcriptome-epigenome data of mouse brain to benchmark SPCoral and other methods^31^. Clustering based solely on spatial transcriptome data failed to clearly delineate cortical layers and could not identify the lateral preoptic area (20-lpo); these regions were better resolved by spatial epigenome, which provided complementary chromatin accessibility signatures (**Fig. 3f**). Conversely, epigenome alone failed to adequately resolve the heterogeneity of the cortical plate (cp) region (**Fig. 3f**). The integrated embedding of spatial transcriptome-epigenome was obtained by SPCoral and competing methods (**Fig. 3g**). The spatial continuity of the identified domains was quantitatively evaluated by global Moran’s I (**Fig. 3h**). Seurat_WNN, due to its lack of spatial constraints, got the lowest Moran’s I. The second lowest method was MISO, which requires H&E images as auxiliary input but was not provided in this dataset. In contrast, SPCoral, SpatialGlue, and SMOPCA exhibited excellent preservation of spatial coherence with higher Moran’s I, and successfully delineated the layered gray matter structures across different cortical layers. Notably, SPCoral and SpatialGlue further resolved the cp region into fine-grained substructures (3-cp and 12-cp) as observed in the transcriptome.

Furthermore, we validated SPCoral’s cross-resolution integration capability. We down-sampled the spatial transcriptome data into lower resolution (**Fig. 3i, Extended Data Fig. 4a**), and used SPCoral to integrate it with the original spatial epigenome data. The visualizations of integrated embedding clearly demonstrated that the cross-resolution integration successfully delineated the fine-grained brain structures at the two distinct resolutions (**Fig. 3j-k**). The Moran’s I for cross-resolution only decreased slightly compared with that for the same-resolution (**Fig. 3h**). Additionally, we used the top-performing methods under the same-resolution condition as controls to assess the cross-resolution accuracy (**Fig. 3l**). The comparison revealed that, despite resolution mismatch and information loss from down-sampled transcriptome data, the clusters of SPCoral’s embedding still presented accurate structures as well as other methods using the same-resolution data. A parallel analysis using down-sampled epigenome data paired with raw transcriptome data yielded analogous conclusions (**Extended Data Fig. 4b, c**), further confirming SPCoral’s robust and reliable performance under substantial resolution differences.

**Fig 4.**
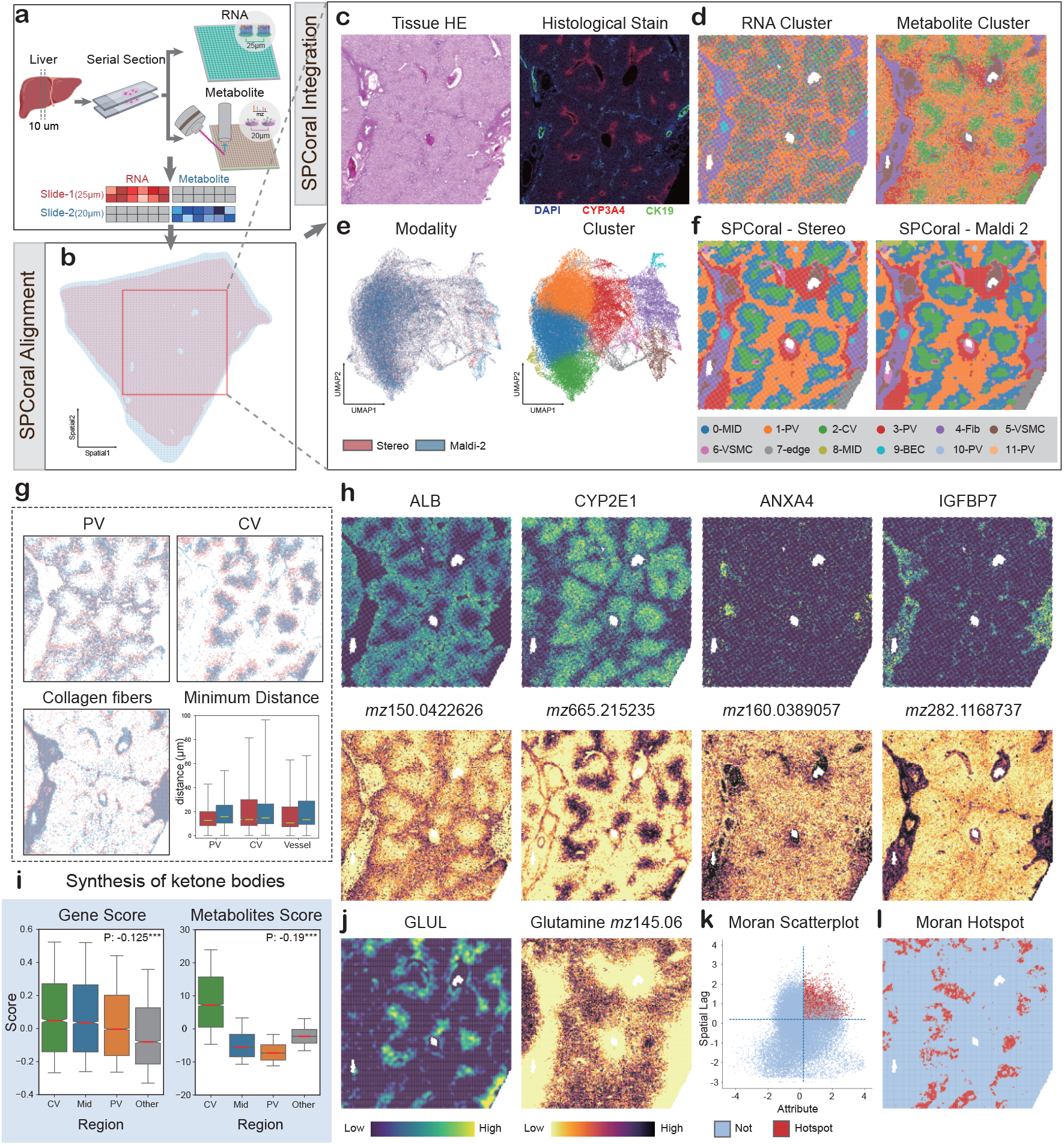
Diagonal integration and downstream analysis of human liver spatial transcriptomics and metabolomics data. **a**, Data preparation workflow. Adjacent tissue sections were profiled using Stereo-seq (transcriptomics at 25 μm resolution) and MALDI-2 (metabolomics at 20 μm resolution). **b**, Spatial alignment results between two modalities. Due to lower quality at the tissue edges in the metabolomics section, analyses were restricted to spots within the red boxed region. **c**, H&E and protein-stained images of the interesting tissue regions. **d**, Independent clustering results within each modality. **e**, UMAP visualizations showing fusion of the two modalities and the identified spatial domains from joint clustering. **f**, In-situ spatial domain maps overlaid on each modality. **g**, Visualization of the interesting regions defined by each modality, overlaid on the aligned coordinate system. Quantification of the minimum distance between same-type spots across modalities. **h**, Feature markers highlighted in each modality. **i**, Bar plots showing enrichment of ketogenesis pathway genes and metabolites across liver zones, with Spearman correlation tests (asterisks indicate P < 0.0001). **j**, In situ expression maps of GLUL (glutamine synthetase) and glutamine metabolite. **k**, Bivariate local Moran’s I scatter plot, with x-axis representing GLUL expression and y-axis representing mean glutamine levels in neighboring spots (dashed lines indicate thresholds, both set to 0.1); red points denote statistically significant hot spots (P < 0.05) with values exceeding thresholds. **l**, In situ map of hot spots, highlighting regions of strong co-localization between GLUL and glutamine.

### Feature-independent and cross-resolution integration enables joint analysis of spatial transcriptome-metabolome

Integrating the spatial metabolome with spatial transcriptome has long been one of the most formidable challenges. Unlike direct regulatory relationship between chromatin accessibility (epigenome) and gene expression (transcriptome), the connection between metabolites and RNAs are far less straightforward. Moreover, current biological technologies have not yet enabled simultaneous large-scale detection of metabolites and RNAs in the same tissue slice. Consequently, data obtained from the adjacent slices must undergo spatial integration to address the issues of absence of shared features and difference in resolutions. We used two datasets that incorporated transcriptomics and metabolomics to confirm SPCoral’s ability in diagonal integration. The first dataset consists of two adjacent slices from a human liver: one profiled by spatial transcriptomics (Stereo-seq, 25 μm bin) and an immediately adjacent slice profiled by spatial metabolomics (MALDI-2, 20μm bin) (**Fig 4a**). To reduce memory usage and improve running speed, both omics were down-sampled to a resolution of 100μm (**Extended Data Fig. 5a**) and the resulting alignment matrix was subsequently applied to the original data to achieve high-resolution registration (**Fig 4b**). The most prominent spatial organization in a healthy liver is lobular zonation, which consists of the periportal (PV) zone with bile ducts (marked by CK19 in immunohistochemistry), the pericentral (CV) zone characterized by CYP3A4 expression, and the intermediate zone (**Fig. 4c**). We annotated the spatial locations of PV, CV, and fibrous regions using transcriptomic and metabolomic data, respectively. After SPCoral registration, the annotated regions from both slices were then overlaid in the common registered coordinate space. Visually, the corresponding structures on both slices exhibited excellent spatial overlap (**Fig 4g**). Quantitatively, for each bin in one modality, we calculated the distance to the nearest bin of the same annotated type in the other modality. On average, every bin identified a corresponding bin of the identical structures within 30μm (**Fig 4g**), confirming that SPCoral achieves precise alignment in this tissue.

**Fig 5.**
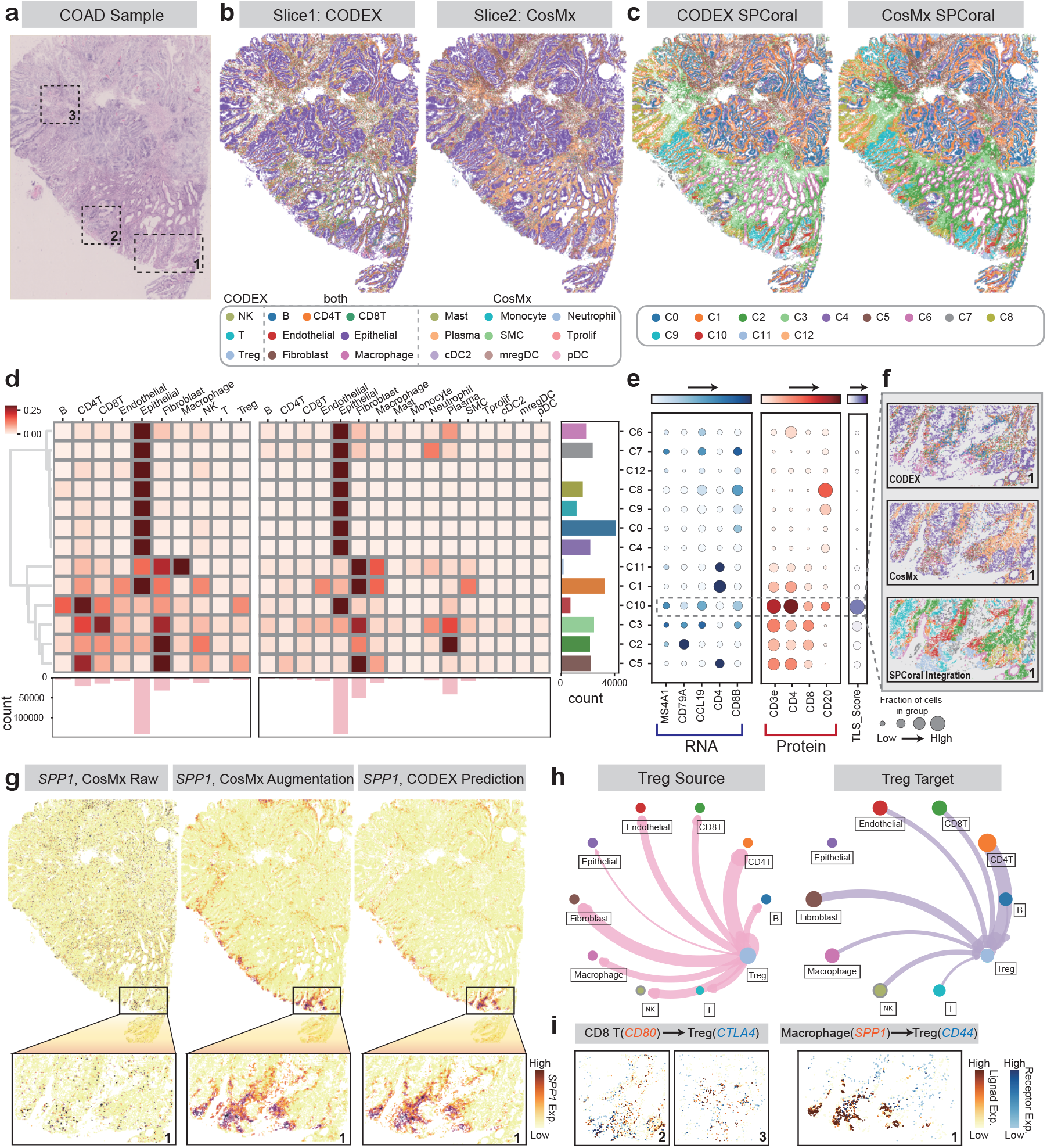
Integration results and applications on COAD multi-omics data. **a**, H&E-stained image of the COAD sample, with dashed boxes indicating three fields of view. **b**, Spatial transcriptomics and proteomics data from the COAD sample. Proteomics data was generated from the first slice using the CODEX platform, and transcriptomics data were acquired from the second slice using the CosMx platform. Colors denote cell types, while the annotations in the dashed boxes represent cell types shared across both omics. **c**, Integration results from SPCoral on this dataset, with colors representing spatial domains identified by joint clustering. **d**, Heatmap showing the correspondence between SPCoral’s spatial domain annotations and original cell-type annotations. Rows are normalized within each omics modality and hierarchically clustered. The bottom bar plot shows cell-type counts, and the right bar plot shows spatial domain counts. **e**, Enrichment of tertiary lymphoid structure (TLS)-related genes, proteins, and their combined scores across spatial domains. **f**, Cell types and spatial domain results for the two modalities in Field of View 1, where domain C10 is prominently clustered. **g**, Spatial expression of SPP1. From left to right: the original RNA expression measured by CosMx on the second slice, imputed RNA expression in the second slice via multi-omics integration, and predicted protein expression in the first slice via multi-omics integration. **h**, Cell–cell communication between Treg cells and other cell types. Left: Treg as source cells; right: Treg as target cells. **i**, Co-expression of the representative ligand-receptors in selected Fields of View. Left: Co-expression view of CD80 in CD8+ T cells and CTLA4 in Treg cells. Right: Co-expression view of SPP1 in macrophages and CD44 in Treg cells. Red indicates ligand-expressing cell types; blue indicates receptor-expressing cell types.

Then we performed feature integration of the two aligned slices. To overcome the computational bottleneck caused by large data volumes, we adopted a patch-based training strategy widely used in large image analysis (**Extended Data Fig. 5c**). When clustering was performed separately on each modality, the transcriptomics successful identified bile ducts (9-BEC) but fail to resolve vascular structures (5-VSMC and 6-VSMC). Conversely, the metabolomics data captured most lumens but could not distinguish bile ducts (**Fig. 4d**). Critically, neither modality alone could reliably delineate the three zones of liver lobules. In the joint embedding of SPCoral, UMAP visualization revealed excellent fusion from both modalities (**Fig. 4e**). The integrated clustering not only clearly separated the bile ducts from other lumens but also more clearly distinguished the three distinct zonal layers (**Fig. 4f**). To highlight biological relevance, we identified and visualized marker genes and metabolites in several key clusters (**Fig. 4h and Extended Data Fig. 5d**): the abundances of gene ALB and metabolite m/z 150.04 were higher in the vicinity of PV region; CYP2E1 and m/z 665.21 were enriched in the CV region; ANXA4 could welly mark bile ducts, while metabolite m/z 160.03 distinguished general lumens; and IGFBP7 and metabolite m/z 282.12 were highly expressed in the perivascular fibrous domain. These results underscore SPCoral’s ability to uncover biologically interpretable spatial domains, which are superior to those of single-modality analyses.

Beyond recovering basic spatial structures, the multi-omics integration results of SPCoral can be used for downstream biological analyses. The first application was to compare the activity of metabolite pathways across different spatial structures. Taken the specific ketone body synthesis pathway as an instance, its activity was scored by the expression of key genes and accumulation of metabolites^32^ (**Methods**). Both scores exhibited significant negative correlation with the distance to the CV, and metabolites of ketone bodies were particularly enriched in the CV regions (**Fig. 4i**). This observation is consistent with established physiological knowledge^33^. The second application involved exploring spatial association of gene expression and metabolite abundance. Using the GLUL and glutamine as an example (**Fig. 4j**), they are expected to be associated since GLUL is the rate-limiting enzyme for glutamine synthesis^34^. However, the expression of GLUL and the abundance of glutamine appear to be unrelated in the situ maps (**Fig. 4j**). One possible reason is the noise of individual molecules, especially metabolites, due to dropout and diffusion in spatial omics technologies^35^. Therefore, we utilized SPCoral to reconstruct spatial gene expression profiles onto the metabolome slice. Then the bivariate Moran analysis between GLUL and glutamine identified statistically significant hotspots (**Fig. 4k**), which indicated the spatial locations where the GLUL-mediated glutamine synthesis preferentially occurred (**Fig. 4l**). These results demonstrated that SPCoral’s diagonal integration and cross-modal prediction are crucial for improving the capability of cross-modal analysis.

The second dataset is derived from a mouse brain, consisting of one section profiled by 10X Visium spatial transcriptomics and an adjacent section profiled by MALDI-TOF spatial metabolomics^36^ (**Extended Data Fig. 6a**). SPCoral’s alignment module successfully registered the two adjacent slices despite the complete absence of shared features and partial tissue overlap (**Extended Data Fig. 6b**). After this alignment, the integration module generated spatially continuous domains that overcome the limitations of the single modality (**Extended Data Fig. 6c-e**): the transcriptomics data alone failed to resolve cluster C8, while the metabolomics data could not reliable distinguish C0 and C2 (**Extended Data Fig. 6f**). The spatially domain-specific features corresponded to genes or metabolites, which embodied the complementarity between the two omics. For example, the metabolomics-missed domains C0 and C2 were strongly characterized by the gene expression of Penk and Nnat, respectively; whereas the metabolites m/z 862.6 and m/z 346.81 specifically highlighted C6 and the metabolomics-exclusive C8, respectively (**Extended Data Fig. 6g, h**). These two datasets proved that diagonal integration of SPCoral is robust for different spatial technologies and resolutions.

### Single-cell spatial integration better decodes immune environment

In the final example, we highlight SPCoral’s capability in handling large-scale single-cell-resolved spatial datasets (two adjacent slices contain over 550, 000 cells) and the downstream analyses of the immune environment (**Fig 5a**). We utilized spatial proteome data (CODEX with 16 marker proteins) and transcriptome data (CosMx 6K gene panel) of adjacent slices from the colorectal adenocarcinoma (COAD) sample^2^ (**Fig 5b**). Proteomics enables precise identification of specific proteins and immune cell types, but it cannot study proteins outside the panel; Transcriptomics provides a broader capture of genes, but the dropout significantly reduces the detection sensitivity of lowly expressed genes, thereby compromising the accuracy of subsequent analyses^37^. These limitations of single omics could be addressed via multi-omics integration and cross-modal predictions of SPCoral.

Firstly, we defined 13 spatial domains (C0-C12) using the integrated embedding (**Fig 5c**) and compared the identified spatial domains with cell-type annotations derived independently from each omics (**Fig 5d**). Some domains were predominantly consisted of epithelial cells; Domains C1 and C11 were primarily composed of fibroblasts and macrophages; C2, C3, and C5 contained fibroblasts mixed with other immune cell types; C10 was particularly noteworthy, appearing as a heterogeneous immune cell cluster in the proteome data while being dominated by epithelial cells in the transcriptome data (**Fig 5d**). The high proportion of CD4 T, B and CD8 T cells in the proteome data suggests C10 may be a tertiary lymphoid structure (TLS). However, there is no obvious concentration of TLS-related markers^38,39^ and TLS score in the C10 region of spatial transcriptome (**Extended Data Fig. 7a, c**). To address this, we used SPCoral to predict protein expression in the transcriptomic slice (**Extended Data Fig. 7b**), and subsequently combined TLS markers from both omics for joint scoring. C10 exhibited a significantly higher TLS score compared to all other domains (**Fig 5e-f and Extended Data Fig. 7d**).

Secondly, we explored the effects of data augmentation by SPCoral. Taking SPP1 as an example, whose expression was barely detected in spatial transcriptome (**Fig. 5g left**). After SPCoral integration, its expression can be augmented by intra-modal imputation (**Fig. 5g middle**) or cross-modal prediction (**Fig. 5g right**). Previous study reported SPP1 majorly expressed in macrophages^40^. The augmented expression profile showed SPP1+ cells exhibited more pronounced expression co-localization with CD68+ macrophages (**Fig. 5g**), which better aligned with the prior study. Although SPP1 was not well detected in transcriptome and not included in the panel of proteome, SPCoral learned its expression pattern through the inherent connections between proteome and transcriptome.

Finally, we demonstrated the advantages of multi-omics integration for cell-cell communication analysis. The regulatory T cells (Treg) was annotated by proteome data via the protein marker CD3e and FOXP3, but Treg was not resolvable from the transcriptome data (**Fig. 5b**). Therefore, it was impossible for any single omics data to directly analyze the ligand-report interactions of Treg with other cells. After multi-omics integration and prediction using SPCoral, Treg cells, ligands and receptors were successfully mapped within a single slice, which made the cell communication analysis possible. We applied the spatial omics tool SOAPy to predict potential ligand–receptor pairs mediating these interactions^41^. Overall, Treg cells showed strong interactions with fibroblasts and other immune cells, but relatively weak interactions with epithelial cells (**Fig. 5h, Extended Data Fig. 8a, b**). Two representative ligand-receptor pairs were CD8-CTLA4 and SPP1-CD44. Treg cells expressed CTLA4 to receive signals from CD8+ T cells^42^, and similarly expressed CD44 to receive secreted SPP1 signals from macrophages^43^ (**Fig 5i**). Both of these communication axes have been implicated in tumor immunosuppressio^44,45^. Taken together, SPCoral enables single-cell level spatial multi-omics integration, significantly enhances downstream analyses, such as spatial clustering, the characterization of expression patterns, and the inference of cell communication.

## Discussion

Integration of spatial multi-omics data is a critical task for understanding spatial microenvironment. Although transcriptome-centered integration methods have made progress in recent years, there are challenges in generalizing to additional modality (e.g., metabolism), or more diverse resolutions, or large-scale single-cell spatial omics. We sought to meet these needs by developing SPCoral. Its core innovation and advantage lie in enabling cross-slice integration without requiring shared features or assumption of consistent spatial resolution.

SPCoral was evaluated across multiple simulated and real datasets encompassing transcriptomics, epigenomics, metabolomics, and proteomics generated by diverse sequencing platforms, with resolutions ranging from multi-cellular spots to the single-cell, demonstrating its reliability and broader applicability compared to competitors. The application on three real-world datasets further confirmed SPCoral’s power through downstream analyses, including spatial domain delineation, cross-modal expression imputation and prediction, spatial correlation assessment, and cell–cell communication inference—highlighting how multi-omics diagonal integration enables more comprehensive and biologically interpretable spatial cell–cell communication inference than single-modality approaches. To overcome computational constraints, we implemented down-sampling and patch-based partitioning strategies, enabling efficient processing of ultra-large dataset, such as two single-cell-resolution slices exceeding 550,000 cells in total, on a single NVIDIA RTX 4090 GPU with 24 GB memory. These strategies do not affect the accuracy of subsequent alignment and integration.

Nevertheless, we acknowledge certain limitations of SPCoral: SPCoral is designed for two adjacent slices, future iterations of SPCoral may need simultaneous integration of multiple adjacent slices; The PCC-based extraction of shared spatial patterns may not be universally effective across all datasets; Performance may degrade when the differences of spatial resolution are extreme. Looking ahead, we plan to extend and improve SPCoral in multiple directions. First, exploring more robust representations of spatial multi-omics patterns to strengthen alignment in the absence of shared features. Second, incorporated histology images or prior knowledge graph to facilitate more efficient and interpretable integration. Third, incorporating joint learning across multiple samples to obtain richer, less biased representations. Finally, applying these advances to more accurate integration and multi-modal prediction.

As research on biological systems increasingly incorporates multiple spatial technologies, SPCoral will play critical roles in integrating any two distinct spatial omics, with no limitations on technology and resolution. We envision that SPCoral can uncover hidden biological insights and facilitate the understanding of complex tissue organization.

## Methods

### Model Structure

#### Alignment model

##### Input

The input to the alignment module consists of three main components. First, the expression matrices of the two omics are provided: 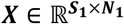 and 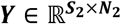, where **S** denotes the number of spots or cells and **N** represents the number of molecules. Second, the respective spatial graphs for each omics are included: 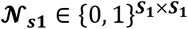 and 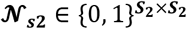, which are constructed using either a k-nearest neighbors (KNN) approach or a fixed distance radius threshold based on spatial coordinates. Third, optionally, shared-feature expression matrices are incorporated 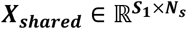 and 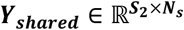, where ***N***_***s***_ is the number of shared features.

##### Feature extraction used GAT

We construct spatial graphs for each modality and employ two separate double-layer GAT encoders to learn low dimensional embeddings of modality-specific information^46^, taking the modality-specific expression matrices ***X*** and ***Y*** as inputs and the outputs are represented as ***H***_***1***_ and ***H***_***2***_, respectively. If overlapping features exist between the two modalities, an additional shared GAT encoder is employed to simultaneously process the common feature subspace, with the shared-feature matrices ***X***_***shared***_ and ***Y***_***shared***_ as inputs, output embeddings are ***H***_**3**_ and ***H***_**4**_. By these GAT, we obtained four embeddings from three GAT encoders:

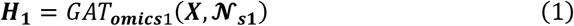

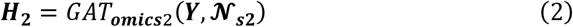

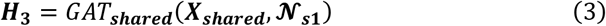

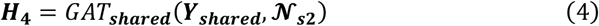

For a detailed description of the GAT computation method, please refer to the Supplementary Information. The decoder architecture mirrors that of the encoder, employing analogous double-layer GAT structures to reconstruct the original expression matrices from the latent representations. The reconstructed matrix is expressed as 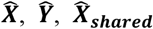 and 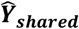.

##### Generation of shared features

For each modality, we construct the spatial adjacency graph **𝒩**_{***1***,***2***}_ as previously described. For every spot/cell ***s*** in the modality, we compute the set of spots/cells at the *i*-th order neighborhood, denoted as 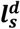 (where *d* represents the neighborhood level). Then we calculate the average expression 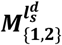 for each molecule across these neighboring spots:

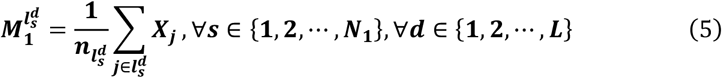

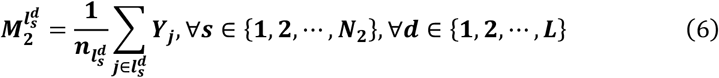

Where 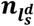 is the number of spots in 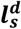, ***L*** is a hyperparameter, representing the maximum number of neighbor layers. For each spot/cell ***s***, we compute the Pearson Correlation Coefficient (PCC) between its expression vector ***X***_***s***_ (***Y***_***s***_) and the average expression vector of its *d* -th order neighbors across multiple neighborhood levels. These PCC values of ***L*** orders are concatenated to form a new spatial covariation matrix ***P***_***1***_ (***P***_***2***_). We posit that the PCC-derived features ***P*** capture modality-agnostic spatial variation patterns and can thus be treated as shared information across the two omics. If the shared-feature embedding ***H***_**3**_ and ***H***_**4**_ is also computed, it is concatenated with ***P***_***1***_ and ***P***_***2***_ to yield an enhanced joint representation for subsequent alignment.

##### Loss function

The loss function consists of two parts: reconstruction loss and adversarial loss.

Reconstruction loss is applied to the original molecules, ensuring faithful recovery of the input expression matrices via the decoder:

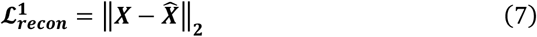

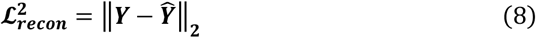

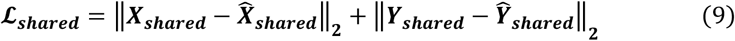

Adversarial discrimination loss is applied to the newly generated features (***P***), where a discriminator ***D*** is trained to distinguish between the two modalities based on these features, while the encoders are optimized to minimize discriminability^47^. The discriminator loss is computed as:

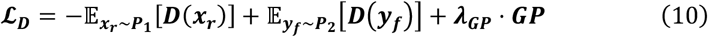

Where ***GP*** is the gradient penalty, ***λ***_***GP***_ is the hyperparameter. After the discriminator updates, we monitor the generator objective as the negative mean critic score on the target anchors:

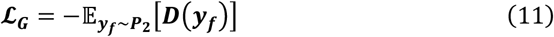

The final loss function is 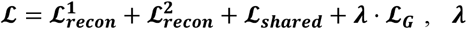 is the hyperparameter and **ℒ**_***shared***_ is optional.

##### Get anchor points from fused Gromov-Wasserstein optimal transport

Subsequently, the learned shared embeddings ***P***_***1***_ and ***P***_***2***_ are fed into an optimal transport framework. We employ the Fused Gromov-Wasserstein optimal transport model, which simultaneously accounts for feature correspondence between anchor points and structural similarity between each spot and its local neighborhood across the two omics^24^:

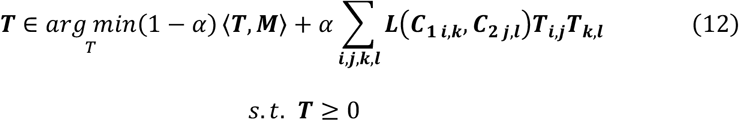

Where the 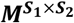 is th<u>e</u> cost matrix between features across omics, it is computed as Euclidean distance between ***P***_***1***_ and ***P***_***2***_ ; ***C***_***1***_ and ***C***_***2***_ are cost matrices in the coordinates space of each omics, ***L*** is a loss function to account for the misfit between the similarity matrices, *α* is trade-off parameter, and ***T*** is optimal transportation matrix.

##### Iterative optimization of anchor points

Based the matrix ***T***, we obtained a set of spot pairs 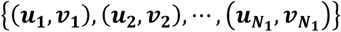, where ***u*** is one spot from omics 1 and ***v*** is the paired spot from omics 2. We calculated the affine transformation matrix from omics 1 to omics 2 using this set. First, we define a 3 × 3 matrix ***A***_***T***_:

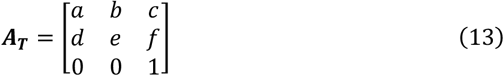

Where *a, b, c, d, e, f* represent the parameters to be determined. Our objective is to apply this transformation to the coordinates of omics 1, thereby minimizing the distance between the transformed coordinates of omics 1 (***U***) and the corresponding anchor points in omics 2 (***V***). It is calculated by the equation:

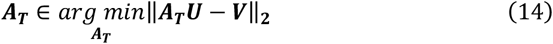

Next, we compute the distribution of distances between each anchor point pair after initial alignment. Anchor pairs with post-registration distances exceeding a threshold (by default, beyond two standard deviations from the mean) are removed as outliers. The affine transformation matrix ***A***_***T***_ is then recomputed using the remaining high-confidence anchor pairs. This step is iteratively repeated until no outliers can be further identified, or until the user-defined maximum number of iterations is reached. This iterative refinement process yields an optimized set of anchors and a more accurate global spatial transformation matrix.

#### Integration model

##### Input

The input to the integration module comprises the following components: the expression matrices of the two modalities, 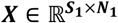 and 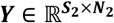 the respective spatial graphs for each modality, 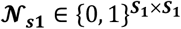 and 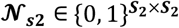 the respective feature similarity graphs for each modality, 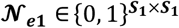 and 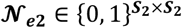, built using a KNN approach based on expression profile similarity; and the joint spatial graph between the two omics layers, 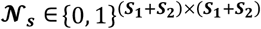, constructed using the Shared Nearest Neighbors (SNN) algorithm on the aligned spatial coordinates^48^.

##### Encoders

First, we process each modality independently. For each modality, the expression features are concatenated with two graphs and fed into a shared-parameter GAT encoder. This encoder generates two distinct embeddings: one based on the spatial graph and the other based on the expression similarity graph, denoted as ***H***_*s*_ (spatial embedding) and ***H***_***e***_ (expression embedding) for that modality. The final outputs of the encoders are denoted as ***H***_*s*1_, ***H***_*e*1_, ***H***_*s*2_, and ***H***_*e*2_. The message passing of GAT is as follows:

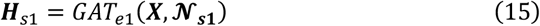

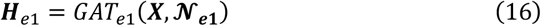

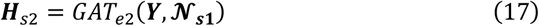

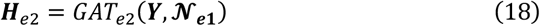

We combine these representations into ***H***_1_ and ***H***_2_ using the following formulation:

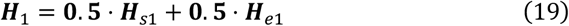

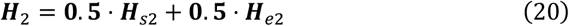

##### Cross-modality attention

Cross-modality attention learning is the core of SPCoral’s integration module, enabling effective information fusion between the two modalities^27^. We employ two independent double-layer MLP networks to extract information from ***H***_1_ and ***H***_2_, respectively. For omics 1, two tensors of identical shape, ***Q*** (query) and ***V***_***1***_ (value), are generated. Similarly, for omics 2, two tensors ***K*** (key) and ***V***_***2***_ (value) of the same shape are produced. The computations are defined as follows:

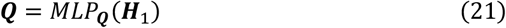

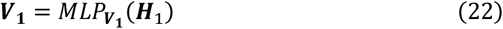

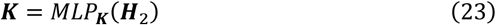

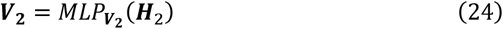

Next, we compute the cross-modality attention matrix **𝒲** using the ***Q*** and ***K*** as follows:

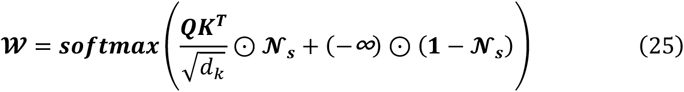

Where *d*_*k*_ represents the column dimension of ***Q*** and ***K*** . Through this attention matrix, we achieve information fusion between the two modalities. Specifically, the updated representation for one omics is obtained by attending to information from the other omics:

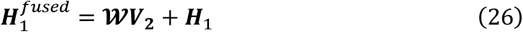

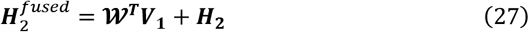

with a residual connection to preserve original modality-specific signals and stabilize training. This bidirectional cross-modality attention can be stacked across multiple layers to progressively enhance mutual information exchange.

Finally, to promote further information exchange between the two modalities, we apply an additional GAT layer. This GAT takes the fused representations 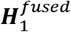 and 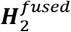 as input node features and propagates information across the omics defined by **𝒩**_***s***_. The GAT update yields refined embeddings:

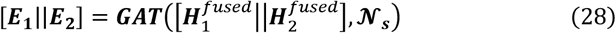

***E***_***1***_ and ***E***_***2***_ represent the final embeddings produced by SPCoral, each incorporating complementary information from both modalities.

##### Decoders

The decoders in SPCoral consist of two GATs, each of which is dedicated to reconstructing the features of one omics modality. By passing ***E***_***1***_ and ***E***_***2***_ through these decoders, we obtain both the reconstruction 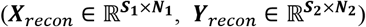 of the original expression profiles and cross-omics predictions 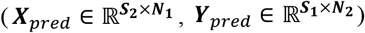 for the features of one modality in the coordinate system of the adjacent slice. The computations are defined as follows:

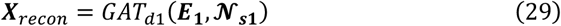

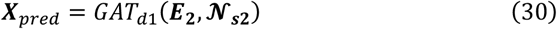

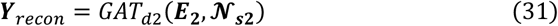

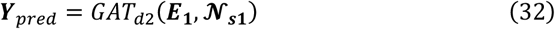

##### Loss function

The loss function **ℒ**_***recon***_ consists of three components: a reconstruction loss, a correspondence loss, and a spatial loss. Reconstruction loss measures the difference between the reconstructed expression profiles and the original expression profiles, and can be formulated as:

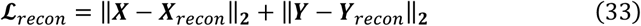

The purpose of the correspondence loss is to enforce complete alignment of information flow across the two omics. To achieve this, we feed the predicted expression profiles back into the corresponding encoders, replacing the original cross-modality attention mechanism, and reconstruct the final embeddings ***E***_***1***_ and ***E***_***2***_:

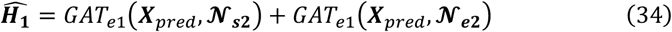

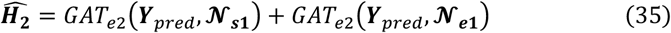

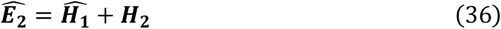

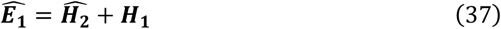

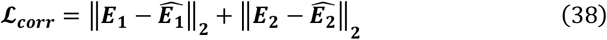

Here *GAT*_*e*1_ and *GAT*_*e*2_ were as same as formulation (15-18).

The spatial loss aims to ensure the spatial continuity of the embeddings ***E***_***1***_ and ***E***_***2***_, thereby enabling downstream analyses to yield well-defined and biologically meaningful spatial domains. It is formulated as:

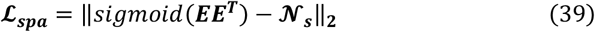

Here, ***E*** refers to the combination of ***E***_**1**_ and ***E***_**2**_. The total loss is defined as **ℒ** = **ℒ**_*recon*_ + **ℒ**_***corr***_ + **ℒ**_***spa***_.

### Benchmarking against competing methods

#### Alignment benchmark

##### Simulated spatial transcriptomics-epigenome data through scMultiSim

To construct datasets suitable for benchmarking against other registration methods, we employed the simulated data generation tool scMultiSim^29^ to produce multiple sets of spatial multi-omics data. Each dataset comprises 130 virtual genes and 390 virtual ATAC peaks, with associations between genes and peaks established via a randomly generated regulatory matrix. Each dataset includes 5 cell types, with spatial positions determined by predefined layouts (enhance, enhance2, island, and layers) and simulated ligand–receptor interactions. Detailed hyperparameters for each dataset are recorded in **Supplementary Table 3**. In the benchmark, we fixed the spatial coordinates of the ATAC modality and applied a random rigid transformation to the coordinates of RNA modality. The performance of each method was evaluated by its ability to recover the original correspondence between the transformed RNA coordinates and the fixed ATAC coordinates.

##### Real spatial transcriptomics-translatomics data of mouse brain

The dataset was obtained from the public data generated by Zeng et al., where the features of the transcriptomic and translatomic profiles exhibit a one-to-one correspondence^49^. To generate simulated datasets with partial feature sharing, we randomly sampled 2000, 100, and 0 features from the original data as shared features, respectively; the remaining features were evenly divided into two parts and assigned to different omics layers, ensuring that this portion of features does not overlap across omics.

##### Method comparison and evaluation

For SPCoral, we first selected 1000 highly variable genes from each omics layer, performed standardization, and then input them into the model. When calculating shared ***P***, the maximum number of neighbor layers was set to 15, and the model was trained for 200 iterations. In the final FGW-OT, the hyperparameter *α* was set to 0.9. For other methods participating in the benchmark, since they do not support the input of unshared features, we first aligned the ATAC features with the RNA features by averaging based on the simulated regulatory network, and then used these aligned features as inputs. We strictly followed the procedures outlined in their official recommended documentation or reference papers, using the default parameter settings provided by each method.

For the simulated datasets, the Euclidean distance between ground truth and post-registration coordinates (*Norm Error*) were calculated for the performance comparison; For the real datasets, we employed four metrics to evaluate the alignment performance of each method (**supplementary information**): the ARI and NMI metrics based on the nearest neighbor regions in the corresponding omics layers, as well as the PCC and COS metrics based on their molecular features.

#### Integration benchmark

##### Simulated spatial data

Following the approach described in SpatialGlue^19^, we utilized code from Townes et al. to generate spatial multi-omics datasets composed of five spatial factors^50^. We simulated two modalities with different data distributions, representing the transcriptome and the proteome, respectively. Omics 1 incorporates a background factor along with two unique spatial factors, while omics 2 includes the background factor and two additional distinct spatial factors. When generating datasets for the two omics, we adjusted parameters and noise levels accordingly.

To benchmark against other methods and to demonstrate SPCoral’s cross-resolution capability, we fixed the size of omics 1 at 1296 spots (36 × 36 grid). For omics 2, we generated data with different spatial resolutions: one with matched resolution (1296 spots, 36 × 36 grid) and another with higher resolution (5184 spots, 72 × 72 grid). The final evaluation metrics consisted of six indices like SpatialGlue^19^: Homogeneity, Mutual Information, V-Measure, AMI, NMI, ARI. For detailed calculation methods, please refer to the supplementary information.

##### Real spatial transcriptomics-epigenomics data of mouse brain

This dataset was obtained from the AtlasXomics platform, providing spatial ATAC-RNA-seq data^31^. Both omics share the same spatial resolution with 20 µm. The transcriptomic profile comprises 22,914 genes, while the epigenomic profile (ATAC) consists of 121,068 peaks. To evaluate SPCoral’s ability to integrate multi-omics data from different resolutions, we constructed the down-sampled datasets of both the transcriptome and epigenome. At the original resolution, every four adjacent spots were aggregated to form one new spot in the down-sampled dataset (**Extended Data Fig. 4a)**. For the transcriptome data, we selected 3000 highly variable genes, performed normalization, and then applied Principal Component Analysis (PCA) for dimensionality reduction, retaining 50 principal components. For the epigenome data, we retained 3000 highly variable features and used Latent Semantic Indexing (LSI) for dimensionality reduction to 50 dimensions. The spatial network was constructed using the first-order neighbors and input into each omics. Finally, the integrated multi-omics embedding were clustered using the R package mclust with a specified number of clusters set to 20.

#### Optimization for high-resolution and large-scale data

The high-resolution spatial technologies enable the profiling of tens of thousands of cells on a single tissue section, which makes the computation methods always suffer speed and memory issues. SPCoral addresses this problem through two strategies.

### Down-sampling for quicker alignment

In the alignment step, users can generate down-sampled data by specifying the target resolution or the desired number of spots per row and column (**Extended Data Figure 5a**). Features in the resulting dataset are aggregated by either averaging or summation. Next, SPCoral computes the transformation matrix using the down-sampled data; Finally, the transformation matrix is applied to the original data to achieve registration of high-resolution.

#### Patch-partitioning for efficient integration

In the integration step, the entire slice can be divided into multiple smaller and overlapping patches (**Extended Data Figure 5c**). Users specify the number of rows and columns to define the patch shape. SPCoral subsequently filters out patches containing fewer spots based on a predefined threshold. For each patch, an independent spatial graph is constructed, and a weight-sharing model is applied across all patches to learn joint embeddings. After processing, the embeddings from individual patches were seamlessly stitched back to reconstruct the complete high-resolution integrated representation.

### Downstream Analysis

#### Spatial domain

The final embeddings ***E*** obtained from SPCoral can be directly used for clustering to identify spatial domains for each spot. By default, SPCoral employs the Leiden algorithm for clustering. Users can optionally adjust resolution parameters or switch to alternative clustering methods (e.g., Louvain or k-means).

#### Intra-modal interpolation and Cross-modal prediction

The decoder module consists of two distinct components, *GAT*_1_ and *GAT*_2_, designed to reconstruct the features of Omics 1 and Omics 2, respectively. When the latent embeddings (***E***) are fed into their corresponding decoders (i.e., ***E***_***1***_ into *GAT*_1_ or ***E***_***2***_ into *GAT*_2_), the outputs represent the interpolation results of the original omics data. Conversely, by inputting ***E***_***1***_ into *GAT*_2_ or ***E***_***2***_ into *GAT*_1_, the resulting expression profiles enable cross-omics prediction.

#### Spatial correlation

Cross-modal expression prediction enables simultaneous access to both modalities’ values at each slice, yielding a co-located and multi-modal feature matrix. This supports spatial correlation analysis using Pearson correlation or global Moran’s I on integrated features, while local Moran’s I identifies cross-modal co-localization regions with coordinated variation or enrichment^51^.

#### Spatial cell communication

Leveraging SPCoral’s cross-modality prediction, data sparsity is significantly alleviated, thereby facilitating the analysis of ligand-receptor mediated cell-cell interactions. We employ the SOAPy toolkit to infer spatially resolved cell–cell interactions^41^. SOAPy provides a permutation-based method that separately models contact-dependent and secreted ligand–receptor interactions. Significant ligand– receptor pairs can be selected using the recommend thresholds of Strength > 2 and Affinity < 0.05.

### Multi-omics datasets processing and analysis

#### Spatial transcriptome-metabolome data of human liver

##### Preprocessing

This dataset was generously provided by the laboratory of Dr. Lijian Hui. Samples were obtained from a normal human liver tissue, which was embedded in gelatin and serially sectioned. The resulting consecutive sections were then subjected to transcriptomic profiling, metabolomic measurement, H&E staining, and immunofluorescence (IF) staining, respectively. Due to noticeable degradation in ionization quality at the edges of the metabolomics (**Extended Data Fig. 5b**), we cropped the raw data by removing peripheral regions and retained only the central, high-quality portion of the tissue for subsequent alignment tests and integration analyses. To enable cell type annotation, the original transcriptomic data from Stereo-seq were binned into 25μm resolution, while the metabolomic data from MALDI-2 were acquired at a native resolution of 20μm per spot.

In the alignment model, we first down-sampled each omics layer and merged them into spots with a resolution of 100 μm. For the transcriptome data, we selected 1000 highly variable genes and performed PCA dimensionality reduction. For the metabolome data, since previous studies have reported that lipid components exhibit greater spatial heterogeneity compared to small-molecule metabolites, we extracted features with larger m/z values, calculated the global Moran’s I index for each feature, and retained those with Moran’s I > 0.2. This ultimately resulted in 155 metabolites selected as input for the alignment model. For both modalities, first-order neighbors were used as edges to construct the spatial networks. The transformation matrix computed from the down-sampled data was then applied back to the original data, yielding the aligned coordinates for the original high-resolution data. In the integration model, we selected 1000 highly variable features from each omics layer and performed PCA dimensionality reduction. We divided the original data into 16 patches as the input for the model. Within the model, first-order neighbors were used as edges to construct single-omics networks, while for the cross-omics network, the number of neighbors in the SNN graph was set to 8. Finally, spatial domains were identified using the Louvain clustering method with a resolution parameter of 0.6.

##### Score of the ketone body synthesis pathway

We employed the scoring method from the Scanpy package^52^ to score genes and metabolites associated with ketogenesis. The relevant genes and metabolites were sourced from the KEGG database^32^. The gene set includes HMGCS1, HMGCS2, HMGCL, ACAT2, ACAT1, BDH1, and BDH2, while the metabolites include m/z 101.0244296 (Acetoacetic acid) and m/z 103.0400088 (β-hydroxybutyric acid).

##### Moran hotspots of glutamine production

We calculated the bivariate local Moran’s I index for GLUL and glutamine according to the following formula:

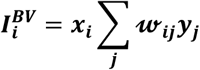

Where ***x***_***i***_ is z-score of GLUL at location ***i***, ***y***_***j***_ is z-score value of glutamine at neighboring location ***j*** . 𝓌_***ij***_ is spatial weight. Hotspot regions were ultimately identified by filtering based on the calculated ***I*** values and the P-values obtained after 1000 permutation tests.

#### Spatial transcriptome-metabolome data of mouse brain

Both transcriptomic and metabolomic data were normalized and subjected to PCA for dimensionality reduction, retaining the top 50 principal components for each modality. In the alignment model, graph edges were constructed by connecting each transcriptomic spot to its first-order hexagonal neighbors, while each metabolomic spot was linked to its four square-lattice neighbors. The number of anchor optimization iterations was set to 10. For the integration model, the intra-modal networks remained consistent with those in the alignment model, with the SNN graph constructed using k=10. Finally, the integrated embeddings were clustered into nine distinct groups using the Leiden algorithm with a resolution of 0.7.

#### Spatial transcriptome-proteome data of human COAD

For the transcriptome data, we performed normalization followed by PCA. For the proteomics data, we applied standard normal standardization (z-scoring) and clipped the maximum absolute value to 5. The original data were divided into 16×16 patches, with each patch containing at least 50 cells in each omics layer. Finally, spatial domains were identified using the Louvain clustering method with a resolution parameter of 1.0. We used the built-in scoring function *score_genes()* in the Scanpy tool to score the TLS gene set. The estimated TLS regions were determined using the upper 5% quantile.

## Data Availability

All datasets used in this study were detailed in Supplementary Table 1. The transcriptomics-translatomics data of mouse brain were downloaded from (https://singlecell.broadinstitute.org/single_cell/study/SCP1835) and the spatial epigenome–transcriptome of mouse brain were from AtlasXplore (https://web.atlasxomics.com/visualization/Fan). The transcriptome and metabolism of mouse brain were obtained from (https://data.mendeley.com/datasets/w7nw4km7xd/1). The transcriptome and proteome datasets from COAD were obtained from SPATCH (https://spatch.pku-genomics.org/#/download). The transcriptomics and metabolomics data from normal human liver tissue were kindly provided by our collaborators and are available after publication.

## Code Availability

An open-source Python implementation of the SPCoral toolkit is accessible at https://pypi.org/project/spcoral. All Jupyter notebooks that produced the findings of the study, including all main and extended figures, are available at https://github.com/LiHongCSBLab/SPCoral. The tutorials of SPCoral are available at https://spcoral.readthedocs.io/en/latest/.

## Author Information

### Authors and Affiliations

Shanghai Institute of Nutrition and Health, University of Chinese Academy of Sciences, Chinese Academy of Sciences, Shanghai, 200031, China.

Heqi Wang, Jiao Yuan, Kunshi Li, Xufeng Chen, Xi Yan, Ping Lin, Zhixuan Tang & Hong Li

State Key Laboratory of Cell Biology, CAS Center for Excellence in Molecular Cell Science, Shanghai Institute of Biochemistry and Cell Biology, University of Chinese Academy of Sciences, Chinese Academy of Sciences, Shanghai, China

Baihua Wu, Haitao Nan, Lijian Hui

State Key Laboratory of Genome and Multi-omics Technologies, BGI Research, Shenzhen 518083, China

Yiwei Lai, Yuan Lv, Miguel A. Esteban

Shanghai-MOST Key Laboratory of Health and Disease Genomics, Shanghai Institute for Biomedical and Pharmaceutical Technologies, Shanghai 200237, China

Lu Xie

School of Public Health, Fudan University, Shanghai 200032, China

Lu Xie

Children’s Hospital of Fudan University and the Shanghai Key Laboratory of Medical Epigenetics, Institutes of Biomedical Sciences, Fudan University, Shanghai, China.

Gangqi Wang

### Contributions

H.L. and H.W. proposed the concept and designed the project. H.W. implemented the code of SPCoral and wrote the manuscript. H.L. supervised the project. J.Y., K.L., and X.C. provided suggestions for the construction of the model. X.Y., P.L., B.W., H.N. and G.W. provided suggestions for the biological explanations of the examples. Y.L., Y.L., L.H. and M.E. provided the spatial transcriptomics-metabolomics data of a liver sample. H.L., L.X. and Z.T. reviewed and corrected the manuscript.

## Acknowledgements

This work was supported by Noncommunicable Chronic Diseases-National Science and Technology Major Project (2024ZD0531300), National Natural Science Foundation of China (32470707, 32300555) and Shanghai Municipal Science and Technology Major Project.

## Extended Data

**Extended Data Fig 1. Additional simulated data for alignment evaluation**

Generated using scMultiSim, showcasing four additional layouts beyond the main figures. **d**, layers layout. For each layout, four additional simulated datasets are presented. Each dataset displays (from left to right): in situ map of ground-truth cell types, pre-alignment spatial positions of the two modalities, and alignment results obtained by SPCoral.

**Extended Data Fig 2. Anchor point visualization during the alignment process**

**a**, Distribution of anchor points in the RIBOmap modality corresponding to each annotated region in STARmap. Left column: anchor pairs directly output by FGW-OT without anchor refinement. Right column: anchor pairs after five iterations of refinement and outlier removal.

**b**, Spatial visualization of anchor pairs, with 300 randomly selected pairs shown. Blue indicates correctly matched regions; red indicates mismatched regions.

**c**, Confusion matrix visualization of the above results, illustrating the correspondence accuracy between regions across paired anchors.

**Extended Data Fig 3. Additional simulated data used for integration evaluation a–b**, Integration results for two additional datasets generated with different random seeds. Each panel includes clustering results for modality 1 and modality 2 at their original matched resolutions, clustering results for modality 2 at mismatched resolution, and integration outcomes from SPCoral and four competing methods (Seurat WNN, SMOPCA, MISO, and SpatialGlue). SPCoral results are presented for both same resolution and different resolution conditions.

**c**, Quantitative evaluation of the above results across six metrics.

**Extended Data Fig 4. Generation and evaluation of cross-resolution mouse brain transcriptomics–epigenomics data**

**a**, Workflow for RNA down-sampling and cross-resolution testing. Every four adjacent spots are aggregated into one new spot. The resulting down-sampled RNA data are paired with full-resolution ATAC data as cross-resolution input to the SPCoral integration model. The integration results obtained at the original resolution are compared with those from other methods evaluated on matched-resolution data.

**b**, Integration results when down-sampling the ATAC data while using full-resolution RNA data.

**c**, Quantitative metrics comparing the original-resolution results from panel b with those obtained by other methods on matched-resolution data.

**Extended Data Fig 5. Additional details for human liver transcriptomics– metabolomics data**.

**a**, Workflow for alignment with down-sampling. Both transcriptomics and metabolomics data are down-sampled to 100 μm resolution. The SPCoral alignment module is then applied to compute the transformation matrix, which is subsequently applied back to the original high-resolution data to obtain alignment at native resolutions.

**b**, Two examples of poor quality at the edges of the metabolomics slice.

**c**, Workflow for patch-based computation. Original omics data are divided into smaller patches based on spatial coordinates for efficient processing, with results subsequently stitched back to the full tissue size. Adjacent patches are configured with partial overlap to ensure seamless reconstruction.

**d**, Find-markers results for spatial domains in the transcriptomics and metabolomics modalities. Markers highlighted in boxes are visualized in Figure **4h**.

**Extended Data Fig 6. Diagonal integration results for mouse brain transcriptomics– metabolomics data**

**a**, Clustering results on the transcriptomics and metabolomics modalities independently.

**b**, Spatial alignment results between the two modalities.

**c**, UMAP visualization colored by modality (distinguishing transcriptomics and metabolomics spots).

**d**, UMAP visualization colored by identified spatial domains from joint integration.

**e**, In situ map of the spatial domain results overlaid on the tissue.

**f**, UMAP visualizations of each original modality separately, colored by the jointly identified spatial domains.

**g**, In situ expression maps of selected feature genes.

**h**, In situ abundance maps of selected feature metabolites.

**Extended Data Fig 7. Additional downstream analysis results for COAD data**

**a**, Spatial expression distribution of tertiary lymphoid structure (TLS) marker genes.

**b**, Spatial expression distribution of TLS marker proteins.

**c**, TLS scores for each spatial domain computed using transcriptomics data alone. **d**, TLS regions identified using transcriptomics-only scoring compared with TLS regions identified using joint transcriptomics and proteomics scoring.

**e**, In situ expression map of the macrophage marker CD68 in the proteomics modality.

**Extended Data Fig 8. Cell–cell communication results involving Treg cells in COAD data**

**a**, Contact-dependent ligand–receptor interactions when Treg cells act as receivers.

**b**, Secreted ligand–receptor interactions when Treg cells act as receivers.

Spot size represents affinity P-value, and color represents interaction strength. Bold borders indicate significant interactions. The ligand–receptor pairs and corresponding cell types highlighted in Figure 5i are marked in red.

## Supplementary information

### Supplementary Information

Supplementary Notes, Methods

**Supplementary Tables 1**

Experimental datasets used in the manuscript.

**Supplementary Tables 2**

Comparison of SPCoral with existing aligned and integrated methods.

**Supplementary Tables 3**

The additional information of benchmark.

**Supplementary Tables 4**

The detailed content of cell-cell communications.

